# Towards a new standard in genomic data privacy: a realization of owner-governance

**DOI:** 10.1101/2024.07.23.604393

**Authors:** Jingcheng Zhang, Yingxuan Ren, Man Ho Au, Ka-Ho Chow, Yekai Zhou, Lei Chen, Yanmin Zhao, Junhao Su, Ruibang Luo

**Affiliations:** Department of Computer Science, The University of Hong Kong, Hong Kong, China; Department of Computing, The Hong Kong Polytechnic University, Hong Kong, China

**Keywords:** genomic data management, data privacy, homomorphic encryption, zero-knowledge proof, blockchain

## Abstract

With the rapid developments in sequencing technologies, individuals now have unprecedented access to their genomic data. However, existing data management systems or protocols are inadequate for protecting privacy, limiting individuals’ control over their genomic information, hindering data sharing, and posing a challenge for biomedical research. To fill the gap, an owner-governed system that fulfills owner authority, lifecycle data encryption, and verifiability at the same time is prompted. In this paper, we realized Governome, an owner-governed data management system designed to empower individuals with absolute control over their genomic data during data sharing. Governome uses a blockchain to manage all transactions and permissions, enabling data owners with dynamic permission management and to be fully informed about every data usage. It uses homomorphic encryption and zero-knowledge proofs to enable genomic data storage and computation in an encrypted and verifiable form for its whole lifecycle. Governome supports genomic analysis tasks, including individual variant query, cohort study, GWAS analysis, and forensics. Query of a variant’s genotype distribution among 2,504 1kGP individuals in Governome can be efficiently completed in under 18 hours on an ordinary server. Governome is an open-source project available at https://github.com/HKU-BAL/Governome.

## Introduction

The advent of affordable advanced sequencing technologies has empowered individuals to explore their health condition through personal genomics, highlighting the critical role genomics plays in modern healthcare ^1^. With increased accessibility, data privacy and security have become an emerging issue when managing personal genomic data. With limitations in storage and analytical capabilities, many individuals opt for third-party services to host and analyze their genomic data. These services offer medical insights and analyses related to ancestry and disease susceptibility, expanding the utility of genomic data beyond the clinical setting.

Despite the benefits, relying on third-party services introduces inherent privacy risks. Users often compromise control over their data by agreeing to terms with limited choices. This leaves the data vulnerable to potential mishandling or misuse, particularly in unregulated contexts. Instances have been documented in which commercial companies share genomic data with pharmaceutical firms in exchange for financial incentives, underscoring the importance of ethical practices related to data security ^2^. Such data mismanagement can have immediate consequences. For example, individuals with high-risk genetic markers are denied life insurance coverage due to undisclosed genomic data usage ^3^. Moreover, when using third-party services, ensuring “The Right to be Forgotten” in the General Data Protection Regulation (GDPR) ^4^, specifically the revocation of data access, is challenging.

The revocation process typically relies on users submitting requests to third parties, who must then comply with relevant regulations. This dependency on third-party compliance makes it difficult to ensure that data access revocation can be executed without undue delay, let alone achieve instant data access control.

The issue of genomic data privacy quickly caught the attention of the academic community, resulting in the development of various methods to protect data. Existing human genomic databases ^5–7^ host research-funded genomic data, and they achieve data privacy by providing access only to successful applicants. Another approach is to provide a unified API for cross-institution genomic data sharing, thereby enabling a centralized gateway with security protocol. Beacon Service, by GA4GH ^8^, was an early attempt at federated data sharing. It aims to achieve collaboration across databases through a distributed storage and sharing network. Despite its intent to facilitate collaboration, the potential for reidentification ^9^ through query analysis remains a critical privacy issue.

Cryptogenomics, which involves applying cryptographic methods to genomic data, is a promising solution for genomic data privacy. Early efforts focused on privacy-preserving data sharing and computation among institutions (also called data custodians). These methods are typically designed for specific genomics analysis tasks, such as cross-institutional single-gene disease diagnosis query ^10^, GWAS ^11,12^ and genetic imputation ^13^. These task-specific protocols by different institutions vary in specializations and capabilities, while none offers personal genome data owner timely and full control of their own data.

Blockchain technology ^14^ offers a new insight to the field of cryptogenomics. Blockchain is a distributed ledger technology that enables multiple participants to engage in secure transactions and information sharing transparently without a central authority, which naturally aligns with the requirements of personal genome data owners retaining full authority over their genomic data without intermediaries as data custodians ^15^. Therefore, starting in 2018, a well-known genomics security contest named iDASH ^16^ extended blockchain to one of its security computing tracks for the task of recording patients’ data sharing consents. There have been attempts ^17^ to store and share genomic data directly using blockchain. While it ensures the security and immutability of transactions, the privacy of information stored on-chain is lost since any participant with read-access to the system can directly access the raw genomic data. Another attempt introduced a citizen-centered method ^18^ that involved both secure computation and a blockchain-based system. However, it supports only simple genomics analysis tasks because only addition operation is supported in its secure computation design, making it impractical for real-life genomics analysis tasks. It also lacks a measurement to avoid data owners and computing parities from providing false information, which is inevitable as the number of participants grows.

We consider that the full-fulfillment of owner-governance is the next step of cryptogenomics. Owner-governance implies three properties throughout the entire lifecycle of genomic data: the data owner retains full authority of her data, 2) the genomic data remains encrypted, and 3) the integrity of both the genomic data and the computation results is algorithmically guaranteed. Practical solutions are urged for a comprehensive owner-governed genomic data management system that should at least include features including user anonymity, dynamic data access control, record audibility, secure data analysis ^19^, and verifiable analysis results. In existing human genomic databases ^5–7,9^, data access revocation is difficult if not undoable once the data has been shared and kept another copy. Queries about data usage logs and permission control are also entirely reliant on the credibility of the data custodian. Thus, establishing a comprehensive system for owner-governed genomic data management is imperative for addressing privacy concerns and empowering individuals in the genomic data landscape.

In this paper, we explored the pathways to achieving the three properties of owner-governance, namely Owner Authority, Lifecycle Data Encryption, and Verifiability. We developed Governome, a realization of owner-governance that fulfills all three properties. Governome utilizes a blockchain to manage all transactions and permissions, enabling data owners with dynamic permission management and to be fully informed about all data usage. It uses homomorphic encryption and zero-knowledge proofs to enable genomic data storage and computation in an encrypted and verifiable form in its complete lifecycle. Data owners can share or unshare their genome in the system instantly. Querying entities can conduct analyses, including individual variant queries, cohort studies, GWAS analyses, and forensics. We benchmarked Governome for different applications and found that querying the population genotype distribution of a random SNP (Single Nucleotide Polymorphism) over 2,504 1kGP ^20^ individuals can be efficiently completed in under 18 hours on an ordinary server. Our experiments demonstrated that Governome can be applied to different genomic data management scenarios at scale. Governome is open-source and available at https://github.com/HKU-BAL/Governome. To our best knowledge, Governome is the first realization of a secure, transparent, decentralized data management system that enables owner-governed genomic data management. We hope that Governome can set a new standard for privacy protection and data sharing in the personal genome era, and in turn benefit personalized medicine and facilitate population genetics researcher at a larger scale.

## Results

### Overview

We defined three properties that lead to the full-fulfillment of owner-governance in a genomic data management system: Owner Authority, Lifecycle Data Encryption, and Verifiability. We developed Governome that fulfilled owner-governance. Governome enables data owners to have 24/7 instantaneous control of their genomic data with full transparency. No plaintext information is stored or generated in the system to eradicate any sort of data leakage. Data integrity and computation result authenticity are algorithmically ensured. Governome supports different genomic tasks, including variant query, cohort study, GWAS analysis, and forensics. We demonstrated Governome’s performance with all variants of the 2,504 1kGP samples, suggesting its robustness when managing large-scale human genome projects and its potential to be scaled-up to managing millions of samples.

### The three properties of owner-governance

We consider a genomic data management system is capable of owner-governance if it simultaneously has the following three properties:

**1) Owner Authority (OA)**: Owners have absolute and instantaneous control over their owned genomic data. At any given time, data owners should be able to modify the access permissions of their genomic data in the system, including revoking data access entirely for any usage. OA also includes data owners’ access to complete data usage logs that are guaranteed to be authentic.
**2) Lifecycle Data Encryption (LDE)**: Data must remain encrypted throughout its lifecycle in the system, ensuring that it is never decrypted or accessed in raw form to protect data security. Encryption should be comprehensively applied to users’ raw data or intermediate computation results in the stage of storage, exchange, and computation. No party, including the data owner, should have direct access to raw information except for the final result provided by the system.
**3) Verifiability (VER)**: Verifiability includes data integrity verifiability and computation process verifiability. Data integrity verifiability refers to the querying entities who initiate a query analysis in the system are able to verify whether the genomic data is free from tampering. Computation process verifiability requires the system to be able to provide evidence for the correctness of the results of any computing process.

### Necessity of the three properties

OA is the core principle of owner-governance, which implies around-the-clock intermediary-free revocation and traceability. Intermediary-free revocation means that the data owners can break away from their previous commitments freely and at any time without any intermediary - they can be the ultimate decision-maker regarding data access or their own data. Traceability means data owners are fully informed, addressing information asymmetry challenges and enhancing control. The combination of decision-making and the right to be informed forms the basis of data owner’s authority over their own data.

LDE is an inevitable requirement for ensuring data security in an owner-governed system. Unless proven otherwise, any disclosure of raw data, even to data owners, will result in potential risks such as information theft and storage device loss, which can have an irreversible impact on data privacy. On the other hand, any party that acquires access to any raw data or intermediate results in plaintext means a deviation from OA since the party can maintain a copy with or without permission, which undermines data owner’s right to decision-making.

VER ensures that the querying entities can always achieve the correct result, which is the foundation of usability. It prevents malicious participants from providing false information that could fake an identity or void research. The use of a blockchain implies crowdsourced data storage and computing. Hence, without a proper mechanism, a dishonest data provider or computing provider might act maliciously and cause permanent damage to the usability of the system. The principle of enabling VER is to trust no one and use mathematical and cryptographic tools to enforce data and computation integrity.

Without OA, data owners would effectively lose control over their genomic data. Without LDE, the genomic data within the system would face inevitable privacy risks when being used. Without VER, the system would loss its trustworthiness, and usability in the end. Therefore, as the next step of cryptogenomics, the three properties OA, LDE, and VER are integral.

### Governome realizes owner-governance

We developed Governome that fulfills the three properties simultaneously. To our best knowledge, it is the first realization of an owner-governance genomic data management system. As shown in Figure 1, Governome includes three layers: 1) a consensus layer to manage agreements among users; 2) a computing layer to manage the different forms of genomic data at various stages, including data storage, exchange, and analysis; and 3) an application layer as an interface for users to interact with the consensus layer and computing layer. The functionality of Governome is built upon the synergy of the three layers. Details about the techniques and design focuses at the three layers are shown in the ‘Feasible approaches to fulfill the three properties of owner-governance’ subsection in Methods.

**Figure 1.**
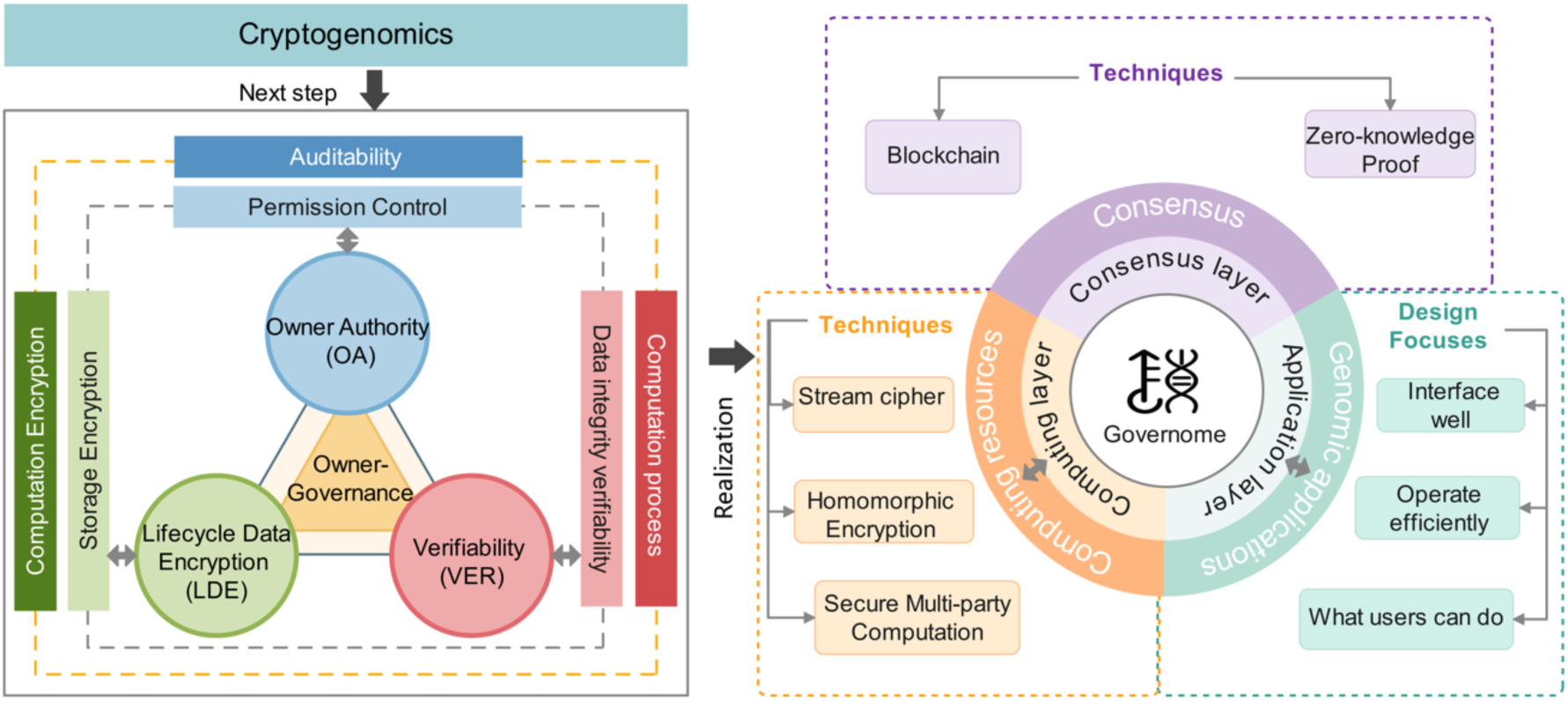
An overview of owner-governance and its realization, Governome. Owner-governance requires three properties, 1) Owner Authority - owners must have absolute and instantaneous control over their owned genomic data; 2) Lifecycle Data Encryption - data must remain encrypted throughout its lifecycle in the system; 3) Verifiability - includes data integrity verifiability and computation process verifiability. Our realization Governome includes three layers that work synergistically, including (1) a consensus layer to manage user agreements; (2) a computing layer for secure computation, and (3) an application layer for genomic applications.

The consensus layer is a blockchain that establishes the ownership of genomic data (see ‘Techniques used at the Consensus Layer’ subsection in Methods). The blockchain stores 1) user permissions settings, 2) metadata and hashes for each query, 3) the source code of supported genomic analysis tasks. Specifically, one’s ownership of her genomic data should be universally recognized, and her modifications to the permissions of her genomic data should not have different versions across different nodes in the blockchain. For each query, (i) the encrypted result, (ii) the individuals involved in serving the query, and (iii) metadata are stored on the blockchain. Owners can achieve auditability by either (a) checking requests that she replied with the access token, or (b) reconstructing the entire logs from the access requests. Moreover, with the support of metadata, hashes and source code, the workflows in Governome are transparent and reproducible by anyone, thus resolving disputes.

The computing layer is for aggregating the storage and computation resources of multiple parties with algorithms (see ‘Techniques used at the Computing Layer’ subsection in Methods). The design of the computing layer focuses on 1) how genomic data is accessed, 2) how the genomic analysis tasks are performed, 3) how multiple parties cooperate to participate in a task. The input of the computing layer is some encrypted data, while the output is fixed-form results of some genomic analysis tasks. Apart from the final output, all intermediate information is computable but cannot be decrypted. The computing layer is responsible for outputting reliable results for tasks, with the computing process being verifiable.

The application layer works as an interface for users who want to use the functions in Governome (see ‘Design focuses at the Application Layer’ subsection in Methods). Considering the steep learning curve of cryptography and secure computation, a user-friendly interface is needed in Governome, while all modules related to privacy and security should be encapsulated within the consensus layer and computing layer. The design of the application layer, on the other hand, focuses on determining who can use Governome and how different users can utilize Governome, where users can simply ask questions predefined by the interface and receive responses. Moreover, as is requested by VER, when users question the reliability of computational results, they should be allowed to request evidence from the interface provided by the application layer and designate someone to verify the data integrity or computation integrity.

### The Workflow of Governome

The workflow of Governome is shown in Figure 2, and the necessary participants in the workflow can be found in the ‘Necessary supporting parties in Governome’ subsection in Methods. To use Governome, a query entity can submit fixed-form queries to the application layer. For example, one can ask, “What’s the genotype distribution of rs6053810 for congenital heart disease patients?”. After checking data owners’ on-chain consent, the consensus layer will send a request to the data owners for an access token (details shown in the ‘How to encrypt genomic data’ subsection in Methods), which can make part of their genomic data accessible to computing layer. After the access tokens for all data owners involved have been collected, the computing layer will pull data from the storage nodes, perform secure computation and return an answer to the query entity through the interface of application layer (details shown in the ‘How computing layer works’ subsection in Methods). Noteworthy, both access tokens and genomic data are utilized in encrypted form. Apart from the fixed-form computation results, no other information is decrypted, thus fulfilling the principle of LDE.

**Figure 2.**
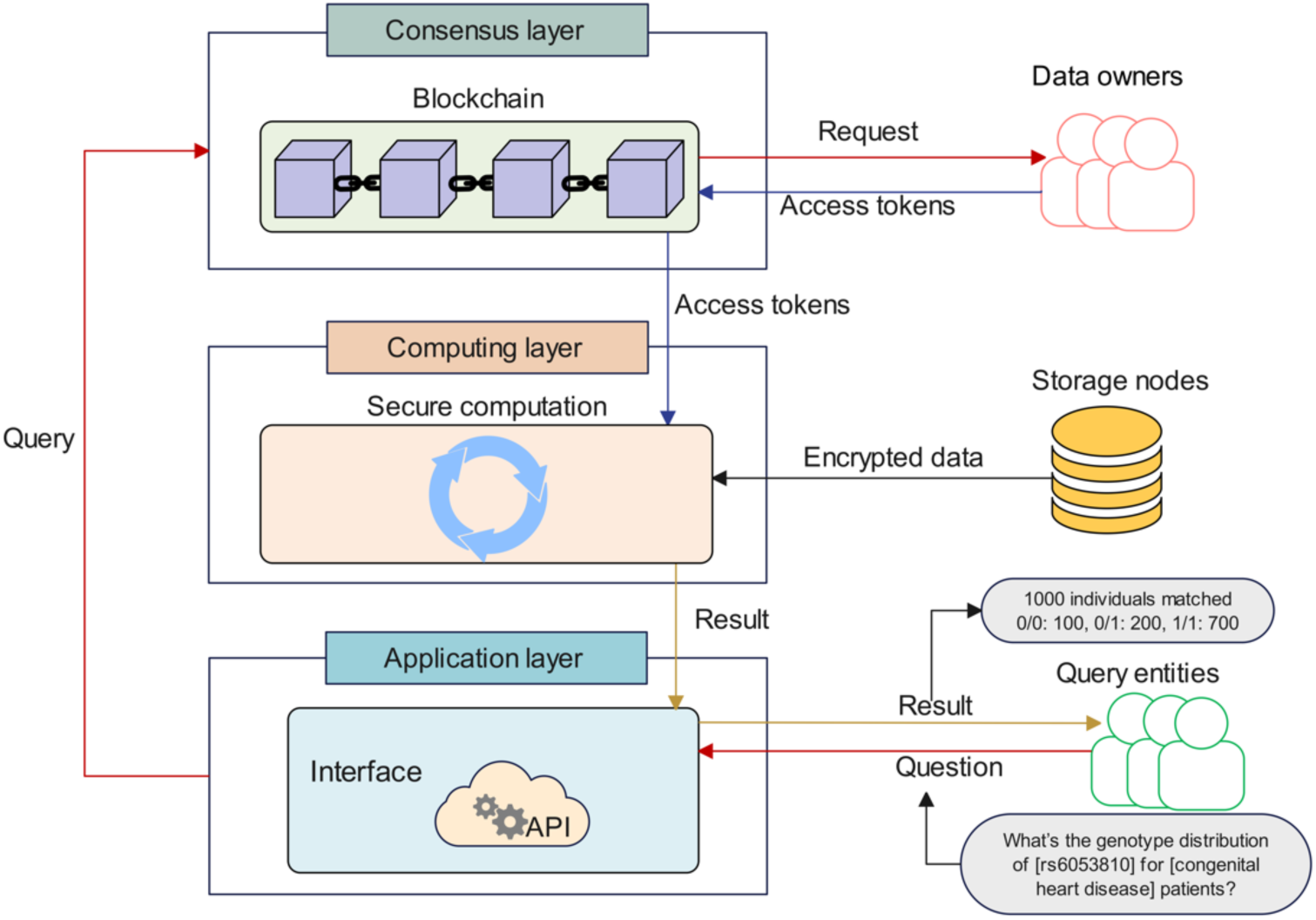
The workflow of Governome. A query entity can ask the application layer a fixed-form question. The application layer will then ask the consensus layer for qualifying data owners. The blockchain managed at the consensus layer will send requests to qualifying data owners, and receive access tokens (See ‘How to encrypt genomic data’ subsection in Methods) from consenting data owners for the downstream homomorphic encryption-based computation. Next, the computing layers will pull the relevant encrypted data blocks of the consenting data owners from storage nodes and perform homomorphic encryption-based computation with the access tokens provided by the consensus layer. No data is decrypted during the computation, except for that the final computation result will be decrypted by the computing layer, and be returned to the query entity with a fixed-form answer.

In general, a data owner is required to be actively responding to requests (specifically, sending access token) from the blockchain, otherwise her data cannot be accessed and would be excluded from analysis. However, it is impractical to require all the data owners to be online and responsive around the clock. Therefore, in Governome, an option is given to data owners to register a precomputed access token so that Governome will skip the data owner and proceed with the token for computing. With this option, a data owner does not need to be active for her data to be used. The registered access token does not need to be recomputed until the next refresh of the computing layer. Details about the precomputed access token can be found in the ‘Precomputed access token’ subsection in Methods.

### Supported genomic analysis tasks in Governome

The application layer has defined a list of genomic analysis tasks, including individual variant query, cohort study, GWAS analysis, and forensics. This section shows the functionalities of the genomic analysis tasks and who can use the functionalities.

Individual variant query allows data owners to explore their own genomic information. Interesting examples including, if someone is interested in whether she suffers alcohol flush reaction after consumption, she can check-up variant rs671 ^21^ that causes aldehyde dehydrogenase 2 deficiency. If a male individual wants to know if he needs to prepare for early-onset hair loss, he can check-up variant rs6152 ^22^ that increases risk of baldness. In Governome, one can input an rsID ^23^ and get the result of her own genotype.

Cohort study allows users to examine the genotype distribution of interested rsIDs relevant to one or more demographics or phenotypes. GWAS analysis allows users to compare a disease cohort against a normal cohort at the interested rsIDs, with p-values returned as results. Cohort study in Governome should obey k-anonymity constraints ^24^. That is, a cohort requires a minimum of k individuals to avoid the risk of being re-identified. The k in Governome is configurable, and Governome returns an error if an analysis forms a cohort with below k individuals. Detailed descriptions of the algorithms used for GWAS are in the ‘HE-based GWAS analysis’ subsection in Supplementary Methods.

Forensics analysis fulfills public security and legal purposes, such as anti-human-trafficking. Given a set of genotypes, Governome can return a list of matching individuals registered in the system. Such an application can bring high social value and is considered to be one of the most important applications of a huge-scale owner-governed genomics database, in addition to research and discovery. However, it is also dangerous, and it compromises personal identity if being misused. Therefore, forensics analysis is exclusive to governmental authorities, and in Governome, we allow a data owner to exclude herself from all forensics analysis, observing our promise to give data owners ultimate control of their data. Forensics analysis can be conducted among all participating individuals in the system, or a smaller group shortlisted by hospitals according to some known demographic characteristics and phenotypes.

Based on the supported genomics analysis tasks available in Governome, we generally distinguished three types of users that demand different analysis permissions (Table 1). The three types are data owners, authorities, and research entities. The ability to perform an individual variant query is exclusively granted to data owners. Using blockchain, data ownership is immediately confirmed, and an individual variant query is processed instantly. In contrast, forensics analysis is exclusive to authorities due to its risk of reidentification. All types of users are allowed to conduct cohort studies in Governome. For a cohort study, if there is a sample list meeting the k-anonymity constraint, the query is processed without further authentication or qualification reviews. The types of users are expandable, and the allowed tasks are configurable in Governome.

**Table 1.** Genomics analysis tasks allowed for different user groups.

|  | Idv. variant query | Cohort study | Forensics |
| --- | --- | --- | --- |
| Data owners | ✓ | ✓ | × |
| Authorities | × | ✓ | ✓ |
| Research entities | × | ✓ | × |

Noteworthy, although Governome has only implemented a few common genomics analysis tasks, it has no limit of having more tasks as long as they can be implemented at the application layer. However, any new tasks need to be sufficiently analyzed and discussed before introducing them into Governome to avoid unintended privacy risks.

### Computational performance of Governome

We evaluated Governome’s computational performance of 1) Generating proofs for an access token, and 2) Homomorphic encryption based computation. These are the two most computationally demanding procedures in Governome. Governome is implemented with programming language Go version 1.21, and all benchmarks were done using the same programming language and version.

#### Generating proof for access token

As mentioned in the workflow of Governome, a data owner should respond to the blockchain and return an access token if consenting the data access request. An access token is the encryption form of an 80-bit key ^25^ kept by a data owner. Apart from the access token, she needs to provide some evidence to show that her 80-bit key is not tampered. Here we have chosen ZK-SNARK (zero-knowledge Succinct Non-interactive Argument of Knowledge) ^26^ as the solution to provide evidence. ZK-SNARK enables a data owner to prove that, without revealing any part of the 80-bit key, 1) she holds the valid 80-bit key according to a hash saved on-chain, and 2) she submitted an access token generated from the valid 80-bit key. More details about why we have chosen ZK-SNARK is given in the ‘Techniques used at the Computing Layer’ subsection in Methods.

The time and memory consumptions are shown in Figure 3. We used a laptop with an Apple M1 CPU and 16GB of RAM, mimicking an average setting of a data owner. The time consumption shows how long it takes to generate a proof and it implies the minimum time a data owner can respond to a data request. The memory consumption shows the peak memory used to generate a proof and it implies how much memory is needed in a data owner’s device in order to respond to a data request. The 80 bits in a key can be used together to generate a proof, or be divided into smaller blocks to generate multiple proofs before merging into a single proof (details given in the ‘Zero-knowledge proof for access token generation’ subsection in Supplementary Methods). The memory consumption increases linear to the block size, but time consumption may vary.

**Figure 3.**
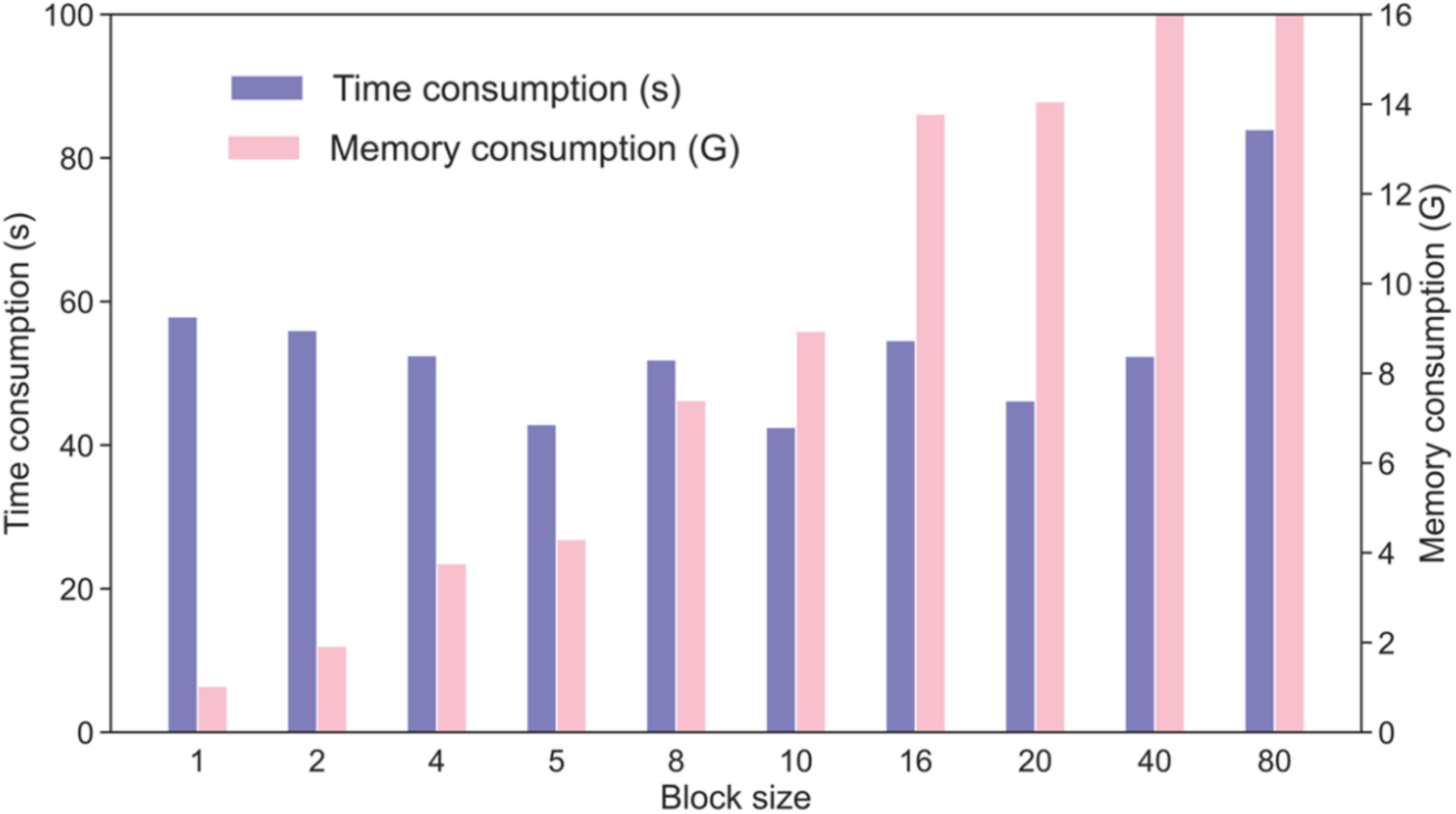
Performance of using ZK-SNARK to generate a proof for an 80-bit key using configurable block sizes ranging from 1 to 80. The memory consumption of block size 40 and 80 exceeded the available memory in our testing device (16GB), and was using memory swap. The exact numbers shown in the figure are given in Supplementary Table 1.

Our benchmark showed that the memory consumption increased from 1.1GB at blocks size 1 to over 16GB at block size 40 or higher. The time consumption varied between block sizes, and had an average of 57 seconds. Since a data owner’s computational capacity is commonly limited to a cell phone or a laptop, a low memory consumption is preferred. We have chosen block size 1 as the default of Governome as it has minimal memory consumption with a moderate time consumption. As a result, a data owner can generate all the necessary information to respond to a data request with a memory consumption of approximately 1GB and time consumption of about a minute, which are completely acceptable.

#### Homomorphic encryption based computation

Homomorphic Encryption (HE) ^27^ is a cryptographic technique that enables computation on encrypted data. In Governome, all data analyses are strictly HE-based computation conducted at the computing layer. Computing nodes at the computing layer are usually powerful servers with many CPU cores and much RAM. For the benchmarks in this section, we used a server with two 32-core Intel Xeon Platinum 8369C 2.9GHz processors and 512GB of RAM. For any genomic analysis tasks, the two computational intensive steps are benchmarked, including: 1) data conversion from stream ciphertext (smaller in size for storage but cannot be used for HE-based computing) to HE ciphertext (larger in size for HE-based computing (details given in the ‘Computation setup in Governome’ subsection in Methods), and 2) data analysis that uses HE-based computation. For samples, we used the 1000 Genomes Project (1kGP) dataset ^20^ comprising the whole genome variants of 2,504 individuals. The mock-up phenotypes of the 2,504 individuals were provided by the Hail library and are available from its tutorial ^28^. The extracted phenotypes were already normalized as either binary or categorical variables. We divided the samples into five cohorts for our benchmarks according to the five superpopulations: Africans (AFR), Admixed Americans (AMR), East Asians (EAS), Europeans (EUR), and South Asians (SAS) defined in 1kGP.

##### Individual variant query and cohort study

Individual variant query is the simplest task in Governome. Our benchmark showed that using a single CPU core, querying any random variant in an individual used at most 15 minutes to return a result. Cohort study, in comparison, demands much more computations, especially when the cohort size is large, and when GWAS analysis is needed. In cohort study, Governome allows inputting rsIDs to specify the variant of interests, and demographic characteristics and phenotypes for choosing samples. If a cohort study query generates no error, Governome will return the number of chosen samples, and the genotypes ratio of the chosen samples at each rsID. The performance of querying a variant in five cohorts and all 2,504 1kGP samples is shown in Figure 4. Generally, the time consumption of both data conversion and data analysis increased linearly against the number of samples in a cohort. Querying a variant in all 2,504 samples was finished in about 18 hours (13h16m for data conversion and 4h37m for data analysis). More CPU cores can be used for parallel computing when querying more than a variant. The results show that Governome can support any population scale because the linear increase in computation matches the expected linear increase in computing nodes when more individuals are introduced to the system.

**Figure 4.**
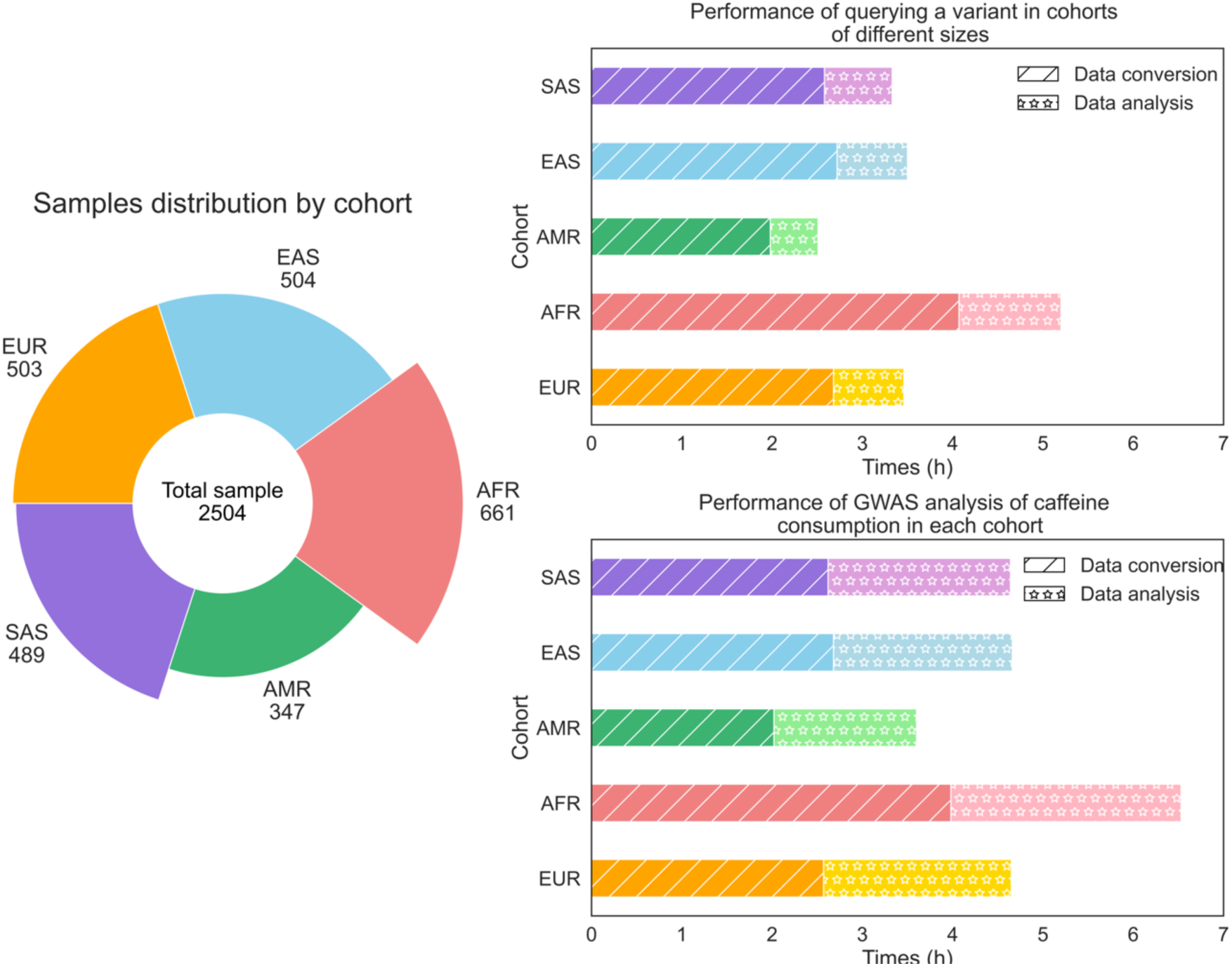
Performance of cohort study, including 1) querying a variant, and 2) GWAS analysis of CaffeineConsumption. The exact numbers shown in the figure are given in Supplementary Table 2 and 3.

For GWAS analysis, a p-value is calculated between the case samples and control samples at each rsID. We have chosen the phenotype ‘CaffeineConsumption’ to divide the samples in each cohort into case (CaffeineConsumption > 4) and control (CaffeineConsumption ≤ 4) samples. The performance of GWAS analysis on rs6053810 is shown in Figure 4. The results have shown that, while the number of samples of each cohort remains the same, data conversion took a similar amount of time, while data analysis took longer due to the algorithmically more complicated p-value computation. Governome allows multiple data analysis tasks to be combined, so data conversion needs to be done just once.

##### Forensics

Forensic genetics relies heavily on analyzing short tandem repeat (STR) loci ^29^. In our benchmark, we have chosen 13 STR loci ^30^ commonly used in forensics for analysis. One can carry out forensics analysis in Governome using cohort analysis at the interested STR loci with all samples in the system included in the cohort. However, the analysis will take excessively long if not impossible to finish when millions of samples are stored in the system. Therefore, we have added an auxiliary data block that stores only the genotype of the 13 STR loci for each individual (see the ‘Auxiliary data block’ subsection in Supplementary Methods). The auxiliary data block is small and specific for forensics analysis. Thus, data conversion can be massively sped up when only forensics analysis is needed. Noteworthy, auxiliary data block can include any number of variants for a specific analysis task in Governome not limited to forensics.

When conducting a forensics analysis, an authority needs to input a list of STR loci with the genotype it is searching for. Additionally, demographic characteristics and phenotypes can be used to reduce the number of samples to be inspected. For each sample, Governome will output a Boolean vector showing a match or mismatch of genotype at each STR loci. As explained in the ‘Necessary supporting parties in Governome’ subsection in Methods, sample IDs in Governome are de-identified. Thus, outside Governome, in order to know the real identity of a matching individual, an authority needs to undergo legal procedures to get a warrant and further work with hospitals.

We tested the performance of the above design on 2,504 1kGP samples. Data conversion took 5 minutes and 51 seconds, while data analysis took 4 minutes. The performance is acceptable if the number of candidates for forensics analysis can be effectively narrowed down by known demographic characteristics and phenotypes.

### Comparing Governome to previous solutions

In this section, we compared Governome against existing genomic data management systems on the three properties of owner-governance. For OA, we extended it into two evaluable dimensions including Permission Control and Auditability. Similarly, LDE was extended into Storage Encryption and Computation Encryption, VER was extended into Data Integrity Verifiability and Computation Process Verifiability. For each system being compared, we assigned either “fulfilled”, “partially fulfilled” or “not fulfilled” for each of the six dimensions, the results are shown in Figure 5.

**Figure 5.**
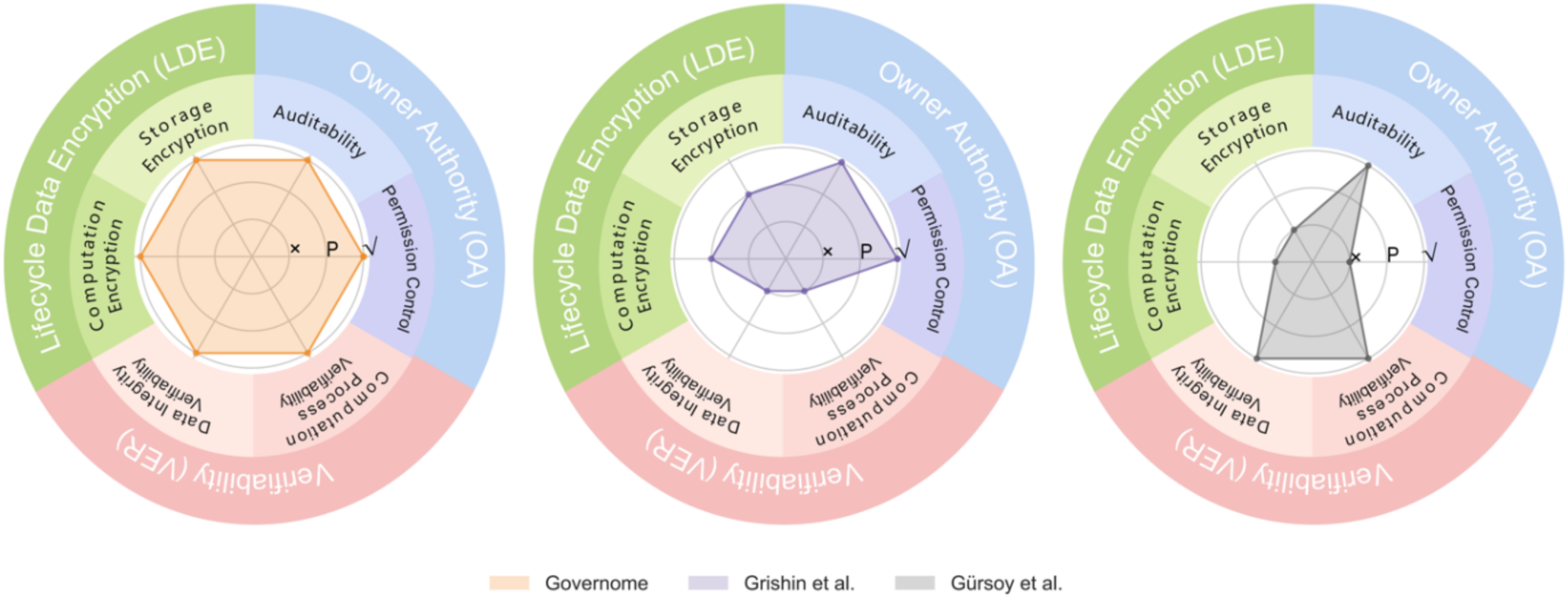
Comparing Governome against existing genomic data management systems on six dimensions including Permission Control, Auditability, Storage Encryption, Computation Encryption, Data Integrity Verifiability, and Computation Process Verifiability. Each dimension was assigned evaluation of either “fulfilled” (✓), “partially fulfilled” (P), or “not fulfilled” (x).

Existing human genomic databases are primarily government-funded centralized databases, such as dbGaP ^5^, UK Biobank ^6^, and AllofUS ^7^, which typically directly avail the data to successful applicants. Some other databases are distributed, but they are still centrally managed and are more like an aggregation of government-funded centralized databases, such as GA4GH beacon ^9^. These databases are centralized and have no capacity of owner-governance, but these are among the most important human genome databases that have promoted the development of genomics in the past decade.

Unlike the traditional human genome databases, Governome is decentralized and owner-governed. There have been similar endeavors that have pushed the field forward, but they are short in one or more dimensions that owner-governance requires.

Grishin et al. ^18^ used a permissioned blockchain that restricts the set of entities that have write-access to the chain, and hence they have achieved full Permission Control and Auditability. They used HE in both computation and storage. But while HE ciphertext is many times larger than the original text in size, they have chosen to only store the HE ciphertext of those who shared their data. This still allows an instantaneously revocation of data access, but resharing data requires uploading the HE ciphertext from the data owner again, which is disincentive to data sharing and an active control of the data permission. The requirement that data owners always need to hold an unencrypted full copy of their data is also what we have avoided in Governome. Thus, we consider Grishin et al. has only partially fulfilled Storage Encryption. In terms of Computation Encryption, we also consider Grishin et al. as partially fulfilled because they used a HE scheme that supports only addition operation, which limited the type and scale of genomic analysis tasks they can support. Grishin et al. has no mechanism to ensure that 1) data owners would not fake their data while uploading, and 2) computing parities would not fake the computing results.

Gürsoy et al. ^17^ used a private blockchain to store BAM (sequencing raw data and alignments) and VCF data without encryption. They achieved Auditability with the use of a blockchain, but since anyone can see everyone’s data on the chain, it puts any sort of Permission Control in vain. They obviously also lack Computation Encryption and Storage Encryption. However, they fulfilled both Data Integrity Verifiability and Computation Process Verifiability because the genomic data is permanently stored on-chain, and any computation can be repeated and verified by others because data is not encrypted.

## Discussion

In this paper, we reviewed the limitations of existing genomic data management systems. We defined the three properties that lead to the full-fulfillment of owner-governance, which is the next step of cryptogenomics. We developed Governome, the first realization of a secure, transparent, decentralized data management system that enables owner-governed genomic data management. With Governome, we demonstrated that the three properties required by owner-governance, including 1) Owner Authority, 2) Lifecycle Data Encryption, and 3) Verifiability, can be fulfilled simultaneously. Governome can do a series of genome data analysis tasks that support the routines of different user groups, including 1) data owners, 2) authorities, and 3) research entities. We benchmarked the performance of Governome and showed its potential to manage large population-scale genomic data.

At the computing layer, Governome uses Torus ^31^, a third-generation homomorphic encryption technique, for homomorphic encryption based computation. To our best knowledge, Governome is the first to use third-generation homomorphic encryption technique for decentralized genomic data management and computing. The second-generation homomorphic encryption techniques used in previous solutions, such as TrustGWAS ^12^, suffer from significant performance degradation when the computing becomes more complicated. This is because the efficiency of second-generation homomorphic encryption relies heavily on single-instruction multiple-data optimization ^32^, which becomes difficult if not impossible when computation becomes complicated and contains excessive branches. Third-generation homomorphic encryption technique has no such limitation, and has enabled Governome to support more complicated genomic data analysis tasks and future expansion.

There are several aspects that could be improved in Governome as future works. First, the current implementation supports only rsID as the variant index. rsID is reference genome agnostic and is verified to be effective and sufficient for personal genome at the stage by public personal genome sequencing services, including 23andMe and Ancestry. However, rsID is incapable of representing every single variant locus of a genome. While more and more personal genomes are now whole-genome sequenced, Governome should support the storage of VCF (Variant Call Format). In fact, Governome can be configured to use VCF to store variants easily. However, the effectiveness of storing the whole genome in personal genomics remains to be debated by the community, especially as we provision that resequencing a genome will get cheaper than storing them permanently.

Governome stores only genomic data and relies on hospitals or institutions that are eligible to host demographic and phenotypic data to shortlist qualifying samples for analysis. It is a practical design considering how most electronic health records are collected and organized. However, technically, Governome can also store and manage demographic and phenotypic data.

Genomic data in Governome is intended to be permanently stored. However, considering the significant advancements in quantum computing, Governome’s security in post-quantum era will become a new challenge. Currently, Governome is not quantum-resistant. In the next step, we will explore optimizing cryptographic methods and privacy protocols to achieve post-quantum reliability.

## Methods

### Feasible approaches to fulfill the three properties of owner-governance

In this section, we discuss the techniques used in Governome and how they serve to fulfill the three properties of owner-governance. As shown in Figure 1, Governome comprises three layers: a consensus layer for authority management, a computing layer for secure computation, and an application layer to make use of the other two layers for genomic applications. At the consensus layer, Blockchain is used to enable dynamic permission control, and Zero-knowledge Proof is used to enforce data integrity. At the computing layer, cryptographic techniques, including Homomorphic Encryption, stream cipher, and Secure Multi-party Computation, are used to fulfill secure computation. At the application layer, several design focuses are introduced to establish the fundamental guidelines for building an efficient owner-governed genomic data management system.

#### Techniques used at the Consensus Layer

#### Why use Blockchain?

Owner Authority strictly requires decentralization, as centralization would technically inevitably jeopardize owners’ control of their data despite how many non-technical promises have been made. Blockchain, as a proven decentralized solution that can achieve consensus, is considered a natural choice for an owner-governed genomic data management system. Blockchain can be configured to ensure that no party can exercise power on others’ data except for their own. Due to the transparent and traceable nature of blockchain, data owner can access their consent and data usage logs at any time, thus ensure auditability. Blockchain can also be used to enforce consensus on computation results so fraud by minorities can be avoided. Both public blockchain and private blockchain are applicable to Governome. While renowned public blockchains are trusted for their decentralization and diversification of users, hence more suitable for publicly or internationally initiated genome hosting, a private blockchain is more flexible and cost-effective for locally initiated genome hosting, in which decentralization is less of a concern.

#### Why use Zero-knowledge Proof?

Lifecycle Data Encryption requires genomic data to remain encrypted throughout its lifecycle in the system. Ciphertext at both storage and computation makes it hard if not impossible to avoid tampering or fraud through traditional means, such as revealing the data or computation results for public scrutiny. In order to ensure the genomic data and computation results are not tampered with or frauded, we use zero-knowledge proof. Zero-knowledge proof ^26^ allows a prover to generate a proof for a proposition without revealing any of its input. In Governome, any data loaded and stored is encrypted with stream cipher, meanwhile a stream cipher key (SCK) is generated and held by the data owner. When a data owner needs to prove she is providing untampered data, Governome uses Zero-knowledge proof to prove that she is providing an encryption of the right SCK to make genomic data accessible without revealing any part of the SCK. Tempered SCK will lead to a different hash that mismatches what has been saved on-chain, thus failing the proof.

#### Techniques used at the Computing Layer

#### Why use Homomorphic Encryption?

The only way to ensure no data leakage is either no data to leak or leakage doesn’t matter. In Governome, LDE mandates that no plaintext exists in the system. Thus, a solution that supports verifiable computation with ciphertext only is needed. Homomorphic Encryption (HE) ^27^ is such a solution that produces deterministic computation results verifiable by all users in the system, using only encrypted data and requiring zero decryption. In contrast, hardware-based solutions, like Intel SGX (Software Guard Extensions) and AMD Memory Encryption Technology, cannot fulfill LDE because they require data to be decrypted when exiting the hardware that supports the same solution. They also cannot fulfill VER because their computational results cannot be easily verified by users who lack the same hardware solution. The details about the HE schemes used in Governome can be found in the ‘Homomorphic Encryption Scheme’ subsection in Supplementary Methods.

#### Why use a stream cipher?

We use HE to fulfill LDE in Governome. However, conversion from plaintext to HE ciphertext (ciphertext capable of HE-based computations) expands the data size by over three orders of magnitude ^31^, making it inefficient if not impossible to store HE ciphertext. In Governome we solve the problem by encrypting plaintext with a cipher that 1) does not significantly increase the size of ciphertext so the ciphertext can be stored efficiently, and 2) can convert from ciphertext to HE ciphertext on-the-fly and without decryption for analysis. Stream cipher ^33^ fulfills these requirements. In Governome, with the use of stream cipher, stream cipher ciphertexts are stored, and will be converted to temporary HE ciphertexts when analysis needs them. More details about how genomic data is encrypted and saved, along with how the stream ciphertext is transferred into HE ciphertext are available in the ‘Storage and computation setup in Governome’ subsection in Methods.

#### Why require multiple parties for the generation of HE key?

Genomic data that are converted into HE ciphertext can be used for analysis, and as the nature of HE, the computation results are also in HE ciphertext. Inevitably, the results are required to be decrypted to become readable before leaving the system. The decryption requires the key that was used to encrypt HE ciphertext. However, that implies that anyone who holds the complete key can decrypt the HE ciphertext to obtain the original genomic data or computation results. In Governome, we 1) required a collaborative generation of a complete key, and 2) avoided any single parties having a copy of the complete key. Our solution uses Threshold Fully Homomorphic Encryption ^34^ (ThFHE), which is a type of Secure Multi-party Computation ^35^ (SMPC). SMPC enables multiple parties to collaborate to generate a key without disclosing the input of any parties, and it ensures the honesty of all parties. Furthermore, SMPC can be used not only for key generation but also for HE ciphertext decryption. So, without revealing the complete key to any party, SMPC then uses the key and the multiple parties who generated the key to decrypt the computation results. The correctness of computation results is ensured by the SMPC protocol ^36–38^. If not using SMPC, one might think of isolation measurements such as limiting the interactions between computing parties (that do compute but are not eligible to see the results) and a HE key holder (that are eligible to see the results), so the computing parties cannot see any intermediate results without the key. However, compliance with such measurements is not algorithmically guaranteed, which is against VER in Governome.

#### Design focuses at the Application Layer

The consensus layer and computing layer together enable owners to have around-the-clock full governance and security of their data. Application layer, on the other hand, is about how to make use of genomic data. An application layer should 1) provide necessary but minimal functions to accomplish different genomic analysis tasks, 2) interface well with the consensus and computing layers, and 3) operate efficiently even with encrypted data. The design of the application layer of Governome has the following major focuses.

#### The application layer defines what the users can do

The application layer defines what could be done with the genomic data stored in the system by providing a set of functions. This set of functions should be meticulously designed to remain necessary but minimal so as to fulfill data management and analysis tasks. The functions are immutable once introduced into the layer so everyone can verify and trust these functions. Availing a function to only a specific set of users enables users to have different roles in the system.

#### The application layer shall do nothing more than the consensus layer and computing layer allow

The application layer uses only the interface the consensus layer and computing layer offered. This allows the applications to have better flexibility, while critical functions such as permission control and result verification are enforced by the consensus and computing layers. For example, data access permission changes received at the application layer will be handled by the consensus layer immediately without a possible delay at the application layer.

#### The application layer needs to work efficiently

Data analysis in Governome works solely with HE ciphertext. A function that works with HE ciphertext needs to be compiled into the combination of single operations like addition over small integer field, the performance of which is expected to be significantly different from, if not much slower than, what the function is supposed to be working with plaintext. Thus, the efficiency of the functions that deal with massive amounts of data at the application layer needs to be carefully examined.

### Necessary supporting parties in Governome

In Governome, besides data owners, multiple parties with different roles are involved to form a robust genomic data analysis system. Their duties and importance are explained as follows.

1. Hospitals or institutions that are eligible to host demographic and phenotypic data: In Governome’s design, it hosts only genotypic data and does not host phenotypic data. This is an effective measurement to 1) isolate different types of critical data, and 2) avoid a single party getting over-powerful. The participating hospitals and institutions connect to the blockchain. Their communications are encrypted and algorithmically verifiable. If a query is asking for a specific cohort with demographic or phenotypic constraints, Governome will ask each participating hospital or institution to provide a list of qualified and anonymous sample IDs. Hospitals and institutions are allowed to return an incomplete (or even empty) list of qualifying samples because they also have the power that equals the Governome to refuse individual data usage. The sample IDs are anonymous by using data owners’ blockchain address.
2. Super users that will never withdraw from the system: Both the consensus layer and computing layer of Governome require multiple active users to maintain functioning. While data owners are granted full governance of their data in Governome, few of them might be active users that can host the blockchain and support the computing in Governome. Super users are a group of users that run servers and can provide storage (i.e., storage nodes) and computing resources (i.e., computing nodes) to keep a Governome system running. Within the data usage lifecycle of genomic data in Governome, super users are responsible for checking hashes and proofs to ensure correctness, pulling data from the storage nodes, converting data into HE ciphertext, and performing the actual computations. The computation results are ultimately decrypted collaboratively by the super users and returned to the query entities through the interface of the application layer. Super users are usually academic institutions, governmental authorities, hospitals, and pharmaceutical companies - the major stakeholders in the system who will mostly benefit from a stable and growing Governome system. In Governome, we require two or more super users to be involved in security-critical procedures, including SMPC, HE key generation, and computation results decryption. While super users have a higher responsibility to keep a Governome system operational, they have data usage privileges identical to all data owners.
3. Temporary computing nodes that temporarily provide additional computing power. Large-scale cohort studies and GWAS analyses are usually conducted by institutional users who are willing to contribute temporary computing nodes in return for some speed up in obtaining a result.

### Storage and computation setup in Governome

#### How to encrypt genomic data

In Governome, raw genomic data is encrypted with stream ciphers and stored in distributed storage nodes that are organized by a blockchain. As data owners have around-the-clock full governance of their data, they are supposed to hold the SCK and are being asked for it every time their data is being used for computing. However, a practical concern is that data owners might leak their SCK due to incidents such as device loss or data theft. The risk is accumulative and gets more significant when the sample size increases. To address the issue, Governome uses hospitals as an additional SCK holder. Instead of holding a complete SCK, data owner and hospital each holds only a part of the key. Governome collects the two partial access tokens (HE ciphertext form of the two partial keys) from the data owner and hospital, and recovers full access token (HE ciphertext form of the complete SCK) with secure computation supported by HE (see Supplementary Figure 1). Details about how genomic data is segmented and stored are described in the ‘Data Segmentation’ subsection in Supplementary Methods, and the discussion of security concerns can be found in the ‘Security of the data blocks’ subsection in Supplementary Methods. This design reduces the risk of leaking the complete SCK. Noteworthy, although hospital also holds part of data owner’s SCK, it has no right over data owner’s genomic data because in Governome, data ownership is ascertained through consensus on the blockchain rather than through the possession of SCK.

#### Precomputed access token

When a data owner’s data is asked to be included for analysis, she is requested to submit an access token generated from her SCK and a Governome-given HE key not only for her data to be used for HE-based computing, but also as a gesture of granting access. However, this behavior requires data owners to respond actively; otherwise, their data would not be included for analysis. This requirement might be too demanding for some data owners who are always willing to be involved in analyses as long as their privacy is protected.

Precomputed access token is such a mechanism in Governome to allow a sharing data owner to register a precomputed access token for accessing all her data blocks (see ‘Data Segmentation’ subsection in Supplementary Methods) so that she does not need to respond to Governome for their data to be used.

#### How computing layer works

The computing layer in Governome involves multiple parties for managing and using stream ciphertext and HE ciphertext, as shown in Supplementary Figure 2. The stream ciphertext from storage nodes will be converted to computable HE ciphertext, using the two partial access tokens collected from the data owner and hospital. The computing layer can carry out different genomic analysis tasks according to the query. The generation of HE key uses SMPC, and therefore no single computing node has the complete copy of it, eliminating the chance that a single computing party could peek or tamper with the HE key. The analysis results are in encrypted form and will be decrypted using SMPC before returning to a query entity.

## Code availability

Governome is available open-source at https://github.com/HKU-BAL/Governome under the BSD 3-Clause license.

## Data availability

The authors declare that all data supporting the findings, including source data and analysis results of this study are available at http://www.bio8.cs.hku.hk/governome/.

## Acknowledgments

R.L. was supported by Hong Kong Research Grants Council grants GRF (17113721) and TRS (T21-705/20-N), the Shenzhen Municipal Government General Program (JCYJ20210324134405015), and the URC fund from HKU.

## Author contributions

R. L. conceived the study. J. Z. and R. L. designed algorithms, implemented Governome. J. Z. and J. S. designed the experiments. R. L., J. Z., J. S., Y. R., M. H. A. and K. C. analyzed the data and drafted the paper. Y. Z., L. C., and Y. Z. evaluated the benchmarking results. All authors reviewed the manuscript.

