## Supplementary Materials for "Towards a new standard in genomic data privacy: a realization of owner-governance"

### Supplementary Figures

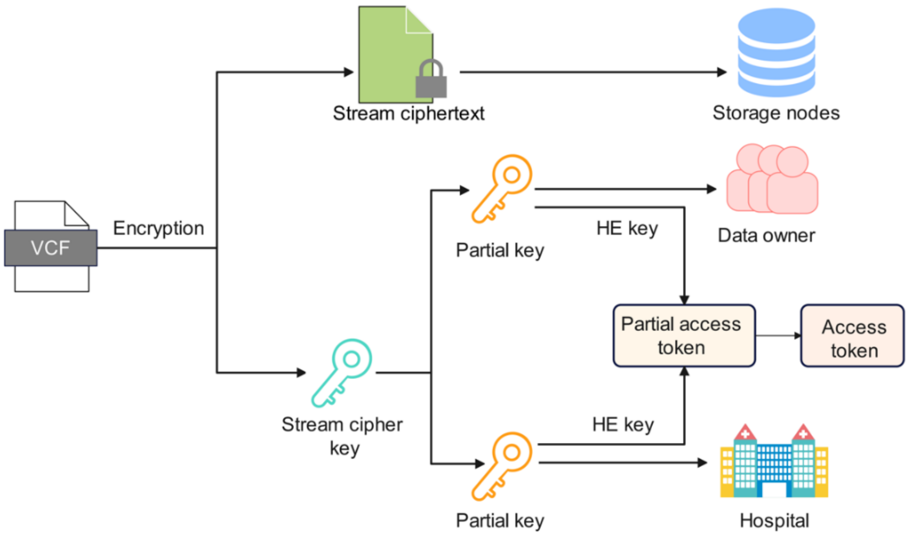

**Supplementary Figure 1. How to encrypt genomic data in our system.**

The storage setup in Governome. Raw genomic data is encrypted by stream cipher and stored as stream ciphertext in rest. Two partial stream cipher keys are kept by the data owner and a hospital, respectively. When being requested, the data owner and hospital encrypt their partial stream cipher key with the HE key into partial access tokens. The two partial access tokens are then being recovered into access token for the genomic data to be convertible from stream ciphertext to HE ciphertext computable in the downstream.

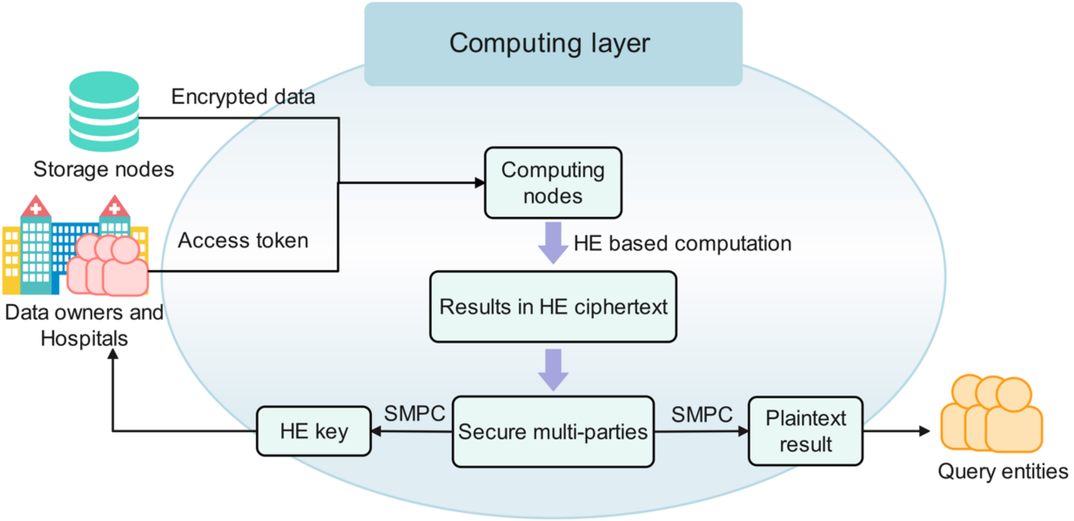

**Supplementary Figure 2. How computing layer works.**

The data flow at the computing layer. The encrypted data and access tokens are sent to computing nodes for HE-based computation. Through secure multi-party computation, encrypted results are decrypted into plaintext result for returned the query entity. All data and information remain in stream ciphertext or HE ciphertext except for the result to be returned to the query entity.

### Supplementary Tables

#### Supplementary Table 1. Performance of using ZK-SNARK generate a proof.

Performance of using ZK-SNARK [1] and Boolean circuit to generate a proof for a 80-bit key using configurable block sizes ranging from 1 to 80. The memory consumption of block size 40 and 80 exceeded the available memory in our testing device (16GB), and was using memory swap.

| Block size | Memory (GB) | Proving Time(s) |
|---|---|---|
| 1 | 1.03 | 57.9 |
| 2 | 1.92 | 56.0 |
| 4 | 3.76 | 52.5 |
| 5 | 4.30 | 42.9 |
| 8 | 7.40 | 51.9 |
| 10 | 8.95 | 42.5 |
| 16 | 13.8 | 54.6 |
| 20 | 14.1 | 43.7 |
| 40 | 23.6 | 61.0 |
| 80 | 43.5 | 110 |

#### Supplementary Table 2. Performance of querying a variant in cohort study.

Time consumption of calculating the genotype distribution of a single variant in different cohorts.

| Cohort | # samples | Data conversion | Data analysis |
|---|---|---|---|
| EUR | 503 | 2h41m | 47m |
| AFR | 661 | 4h4m | 1h8m |
| AMR | 347 | 1h59m | 32m |
| EAS | 504 | 2h43m | 47m |
| SAS | 489 | 2h35m | 45m |
| All | 2,504 | 13h16m | 4h37m |

#### Supplementary Table 3. Performance of GWAS analysis of Caffeine Consumption.

Time consumption of calculating the GWAS analysis *p*-value of Caffeine Consumption in different cohorts.

| Cohort | # samples | Data conversion | Data analysis |
|---|---|---|---|
| EUR | 503 | 2h34m | 2h5m |
| AFR | 661 | 3h59m | 2h33m |
| AMR | 347 | 2h1m | 1h35m |
| EAS | 504 | 2h41m | 1h59m |
| SAS | 489 | 2h37m | 2h1m |

### Supplementary Methods

#### Data Segmentation

Genomic analysis tasks that query variants usually request one or a few variants. Therefore, a data segmentation design that optimizes the query of a small number of variants, while having minimal overhead when retrieving many or all variants is required.

In Governome, we use hash indexing to partition variants into data blocks. The number of data blocks per individual, denoted as *b*, is configurable but unchangeable after system initiation. We suggest storing an average of 40 to 50 variants per data block. For example, an individual with whole-genome sequenced usually has four to five million variants, thus *b*=96k is an optimal setting. When using Governome to store only exome sequencing variants, *b* should be reduced. A principle is that the average number of variants per data block should not go below 20 to avoid data leakage through probing (More details are shown in the subsection 'Security of data blocks'). The storage size of each variant is 36 bits, with 32 bits for rsID and four bits for genotype (16 possibilities including 0|0, 0|1, 0|2, 0|3, 1|0, 1|1, 1|2, 1|3, 2|0, 2|1, 2|2, 2|3, 3|0, 3|1, 3|2, and 3|3). For each variant, the corresponding data block ID is calculated as $\mathrm{SHA3}(sampleID, rsID) \bmod b$. *sampleID* is a string no longer than 15 characters (e.g., "NA12878" in 1kGP). In practice, the *sampleID* can be the user ID of the data owner or her corresponding blockchain address with a random suffix (salt). We used SHA3 as the hash function, which is provided by the Ethereum blockchain as a built-in function that runs efficiently on-chain.

A data block is the smallest data unit to be retrieved for a query in Governome. Each data block is encrypted with a different Stream Cipher Keys (SCK). Each SCK only gives access to a data block. So, multiple SCKs are needed to access multiple or all variants of an individual. This design enables Governome to support fine-grained access control, giving data owners precise control over who can access which data blocks. However, it is not necessary for a data owner to really hold multiple SCKs. In fact, the storage and access of tens of thousands of SCKs is computationally inefficient. In Governome, each data owner holds only a primary SCK. The SCK for each data block is derived from the primary SCK using $MiMC(\text{primary SCK}, segmentID)$. We used MiMC [2] as the hash function here, which is efficient in the ZK-SNARKs circuit.

As elaborated in the 'Storage setup in Governome' subsection of Methods, the SCKs held by data owners and hospitals are partial SCKs. To recover a complete SCK, a data owner and her responsible hospital both need to provide their corresponding partial SCK. A complete SCK is calculated as $\mathrm{SCK}_{\mathrm{dataOwner}} \oplus \mathrm{SCK}_{\mathrm{hospital}}$, where $\mathrm{SCK}_{\mathrm{dataOwner}} = MiMC(\text{primary }\mathrm{SCK}_{\mathrm{dataOwner}}, segmentID)$, and $\mathrm{SCK}_{\mathrm{hospital}} = MiMC(\text{primary }\mathrm{SCK}_{\mathrm{hospital}})$. Notice that hospital uses primary SCK for all data blocks with the rationale that hospital will reject data access to an individual as a whole, instead of rejecting access to particular data blocks.

#### Security of the data blocks

An observation in human genomics is that the number of variants of a normal human is only 0.1-0.2% of the genome size. Thus, as a storage-efficient design, in each data block, only variants with alternative alleles are stored (i.e., variants with reference allele are not stored). However, that enables an attacker to do probing, that is, to inspect the size of a data block to determine the number of variants in the data block. Although the attacker wouldn't be able to decrypt the data block without the correct SCK, it is possible that one can infer whether a variant exists or not in a person if the data block has only one or few variants in a data block and if the assignment of a rsID is the same for each individual.

To address the concerns, firstly, the assignment of a rsID (to which data block) is unique for each individual in Governome by adding salt. Salt is an individual unique value that is added to the input of a hash function to create unique hashes for every input. Governome uses the random suffix in *sampleID* as the individual-unique value.

Secondly, even with a sensible setting of *b*, a small number of data blocks will still contain fewer than 20 variants according to normal distribution. We solved the problem by adding space fillers to a data block to ensure each data block is 20 or more variants long in size. As data blocks are stored in stream ciphertext, attackers wouldn't know how many space-fillers are in a data block. The minimum number of variants per data block, 20, can be raised for better security, but at the price of more wasted storage.

#### Auxiliary data block

Governome allows auxiliary data block besides general data block. Auxiliary data block is task-specific and aims to provide better storage and computation efficiency for specific tasks.

Using forensics analysis as an example. The design of general data block that randomly disperses variants into multiple data blocks with individual-specific hashing implies that retrieving *n* random variants will most likely need to retrieve *n* data blocks. This design was optimized for the best security when the variants of each individual can vary a lot and use only a small subset of all available rsID (otherwise, storing all rsID would be a better solution). However, for forensics analysis that 1) inspects only a fixed set of about tens of variants per individual, and 2) needs to scan over a large number of individuals, the overhead of using general data block becomes significant and caps the scale of Governome. Auxiliary data block addresses the issue by storing the variants required by a task in a single fixed-size data block. Thus, for forensics analysis, only one data block is needed to be retrieved per individual.

In Governome, as a proof of concept, we used an auxiliary data block design that stores 13 commonly used Short Tandem Repeats (STRs) [3], including D3S1358, vWA, FGA, D8S1179, D21S11, D18S51, D5S818, D13S317, D16S539, THO1, TPOX, CSF1PO, and D7S820. The list can be of any size to accommodate different types of tasks. Multiple auxiliary data block designs are allowed in Governome, but the designs are immutable after system initiation.

#### Homomorphic Encryption Scheme

So far, the majority of homomorphic encryption used in bioinformatics practices refers to second-generation homomorphic encryption. As computational complexity increases, the efficiency of second-generation homomorphic encryption technology drastically declines, rendering it impractical in some cases. For instance, while second-generation homomorphic encryption supports GWAS, it is limited to pre-processed data and does not support preprocessing tasks such as data cleaning and quality control. In contrast, third-generation homomorphic encryption does not have these limitations and is considered the best practice for deep neural networks in the era of machine learning. Therefore, we have chosen third-generation homomorphic encryption as the primary technology for the computing layer in Governome. In this section, we review the technical details of the third-generation homomorphic encryption scheme used in Governome.

For better flexibility, in Governome, we chose the third-generation homomorphic encryption scheme TFHE [4], which is based on torus. TFHE computations occur within a small integer domain $T$ and support two types of operations: addition operations and programmable bootstrapping [5]. Addition operations include negation and simple addition, through which we can obtain linear combinations of known ciphertexts. Programmable bootstrapping is a single-variable operation that allows us to construct and implement a single-variable mapping $f: T \to T$, where $f$ can be arbitrarily chosen. In Governome, we construct equivalent Boolean circuits based on these two operations to perform computations.

The hardness of TFHE is based on LWE (Learning With Errors [6]). The encryption process is as follows:

1. $KeyGen: \boldsymbol{s} \leftarrow Z_q^n,\ pk \leftarrow (\boldsymbol{A}, \boldsymbol{b}),\ where\ \boldsymbol{A} \leftarrow Z_q^{n\times n},\ \boldsymbol{b} = \boldsymbol{As} + \boldsymbol{e}$
2. $Encrypt_{pk}(m): ct = (\boldsymbol{a}^*, b^*),\ where\ \boldsymbol{r} \leftarrow \{0,1\}^n, \boldsymbol{a}^* = \boldsymbol{Ar} + \boldsymbol{e_1}, b^* = \langle \boldsymbol{b}, \boldsymbol{r} \rangle + \Delta m + e_2$
3. $Decrypt_s(ct): return\ \boldsymbol{round}(b^* - \langle \boldsymbol{a}^*, \boldsymbol{s} \rangle)$

Additionally, to achieve ThFHE and conceal the HESK (Homomorphic Encryption Secret Key), Secure Multiparty need to collaborate to generate some auxiliary information based on the unseen HESK. This includes: 1) HEPK (Homomorphic Encryption Public Key) for encryption. 2) KSK (Keyswitch Key) and BSK (Bootstrap Key) for computation. 1 will be made public, allowing anyone to encrypt their information using it. 2 will be submitted to the computing nodes; otherwise, the computation cannot be performed.

Noteworthy, the unseen HESK and the aforementioned 1 and 2 need to be updated periodically. Otherwise, even if the data owner revokes sharing, the access tokens they have submitted can always be accessed by attackers.

#### Zero-knowledge proof for access token generation

In order to ensure data integrity, data owners and hospitals need to prove that they have submitted the correct access token without revealing any part of it. To achieve this, we use ZK-SNARK, which is a framework of zero-knowledge proof that allows a prover to generate a proof for a proposition without revealing any of its input. In this section, we will illustrate how an access token and it corresponding zero-knowledge proof is generated.

When a genomic analysis initiates in Governome and the public blockchain asks for access tokens from the corresponding data owners and hospitals, what is collected from the data owners and hospitals are partial access tokens. A partial access token is an encrypted partial SCK encrypted with HEPK, plus an honesty proof to enforce data integrity. The partial SCK is generated from the primary

SCK, whose hash is stored on-chain as a credential. A usual data owner usually uses a smartphone or laptop to interact with Governome. The 'Generating proof for access token' subsection in Results showed that generating a ZK-SNARK proof is a memory demanding procedure. Therefore, we choose to split the partial SCK into smaller blocks to use and reuse smaller zero-knowledge proof circuits that fit smaller blocks to reduce memory consumption. The pseudocode of ZK-SNARKs is shown below, with the use of library gnark [7].

Pseudocode for ZK-SNARKs:

```
    Input:
        Public variables pk, blocksize
        Public witness blockID, segID, credential, ciphertext
        Private witness primarySCK, error
    Procedure:
        mask ← (1 << blocksize) − 1
        block ← hash(primarySCK, segID, blockID) & mask
        assert hash(primarySCK) = credential
        for i from 0 to blocksize − 1 do
            bit ← block & 1
            ct ← LWE(bit, error[i], pk)
            assert error[i] is valid
            assert ct = ciphertext[i]
            block ← block >> 1
        generating a proof π for the assertions
```

As shown in the pseudocode above, we split the partial SCK into $\mathrm{length/blocksize}$ blocks and generate proofs for each block sequentially. The original single proof is divided into $\mathrm{length/blocksize}$ proofs, thus the scale of the proof circuit can be significantly reduced. The "Generating proofs for an access token" subsection in Results shows that when $\mathrm{blocksize}$ is set to 1, the memory consumption is 1.1GB, which can be accommodated by most of the modern laptops.

Moreover, zero-knowledge proof requires even better security regarding the hash function, so instead of using SHA3 that was used in data segmentation and indexing, we used MiMC [2], which is based on large prime fields as the hash function in zero-knowledge proof.

#### HE-based GWAS analysis

In this section, we describe the details about HE-based GWAS analysis that calculates the *p*-value of individual variants against a phenotype using linear regression [8]. Since the algorithm is implemented in a Boolean circuit, division and floating-point operations are very expensive. Therefore, it is necessary to perform transformations on the GWAS formula to convert most floating-point operations in linear regression to integer operations, and to minimize the number of divisions.

To be specific, the linear regression model is defined as follows:

$$\mathrm{y} = \beta_0 + \beta_1 \mathrm{x} + \varepsilon$$

Here, $y$ is the phenotype, while $x$ is the genotype. In our benchmark, we used CaffeineConsumption (1 if CaffeineConsumption>4, 0 otherwise) as $y$, and $x$ as either 0, 1, or 2 (reference, heterozygous variant, homozygous variant).

According to the definition and formula of linear regression, we have:

$$\widehat{\beta_0} = \frac{\Sigma y_i \Sigma x_i^2 - \Sigma x_i \Sigma x_i y_i}{n\Sigma x_i^2 - (\Sigma x_i)^2}$$

$$\widehat{\beta_1} = \frac{n\Sigma x_i y_i - \Sigma x_i \Sigma y_i}{n\Sigma x_i^2 - (\Sigma x_i)^2}$$

To perform a t-test, we need to calculate the t-value as follows:

$$t = \frac{\widehat{\beta_1}}{s_{\widehat{\beta_1}}} \sim t_{n-2}$$

where $s_{\widehat{\beta_1}} = \sqrt{\left(\frac{\Sigma(\widehat{\beta_0}+\widehat{\beta_1}x_i-y_i)^2}{(n-2)\Sigma(x_i-\bar{x})^2}\right)}$

In the above formula, the sample size $n$ is known, and covariance calculation is not needed. Once we calculated the t-value, the p-value can be calculated using the cumulative distribution function.

Furthermore, we can perform the following transformation on the above formula:

$$\frac{n}{n-2} \cdot t^2 = \frac{Numerator}{Denominator},$$

where $Numerator$ and $Denominator$ can be written as:

$Numerator = (n\Sigma x_i y_i - \Sigma x_i \Sigma y_i)^2 (n\Sigma x_i^2 - (\Sigma x_i)^2)$ ,

$Denominator = n(\Sigma x_i y_i)^2(\Sigma x_i)^2 - n(\Sigma x_i^2)^2(\Sigma y_i)^2 - n^2\Sigma x_i^2(\Sigma x_i y_i)^2 + \Sigma x_i^2(\Sigma x_i)^2(\Sigma y_i)^2 + n^2(\Sigma x_i^2)^2\Sigma y_i^2 - 2n\Sigma x_i^2\Sigma y_i^2(\Sigma x_i)^2 + \Sigma y_i^2(\Sigma x_i)^4 + 2n\Sigma x_i^2\Sigma x_i y_i \Sigma x_i \Sigma y_i - 2\Sigma x_i y_i(\Sigma x_i)^3\Sigma y_i$ ,

With the transformation, we can calculate the $Numerator$ and $Denominator$ through HE-based computation. Subsequently, by truncating the $Numerator$ and $Denominator$ to 16-bit precision and performing Goldschmidt division [9], we can obtain an accurate t-value, which is then converted to a p-value in plaintext operations. Notably, while covariance calculation is not needed, the calculation of p-value of each variant is independent, thus can be processed in parallel.
